# Starvation resistant cavefish reveal conserved mechanisms of starvation-induced hepatic lipotoxicity

**DOI:** 10.1101/2024.01.10.574986

**Authors:** Macarena Pozo-Morales, Ansa E Cobham, Cielo Centola, Mary Cathleen McKinney, Peiduo Liu, Camille Perazzolo, Anne Lefort, Frédérick Libert, Hua Bai, Nicolas Rohner, Sumeet Pal Singh

**Author notes:** These authors co-supervised the work and contributed equally. Email for correspondence and. These authors contributed equally (Macarena Pozo-Morales, Ansa E Cobham).

## Abstract

Starvation causes the accumulation of lipid droplets in the liver, a somewhat counterintuitive phenomenon that is nevertheless conserved from flies to humans. Much like fatty liver resulting from overfeeding, hepatic lipid accumulation (steatosis) during undernourishment can lead to lipotoxicity and atrophy of the liver. Here, we found that while surface populations of Astyanax mexicanus undergo this evolutionarily conserved response to starvation, the starvation-resistant cavefish larvae of the same species do not display an accumulation of lipid droplets upon starvation. Moreover, cavefish are resistant to liver atrophy during starvation, providing a unique system to explore strategies for liver protection. Using comparative transcriptomics between zebrafish, surface fish, and cavefish, we identified the fatty acid transporter slc27a2a/fatp2 to be correlated with the development of fatty liver. Pharmacological inhibition of slc27a2a in zebrafish rescues steatosis and atrophy of the liver upon starvation. Further, down-regulation of FATP2 in drosophila larvae inhibits the development of starvation-induced steatosis, suggesting the evolutionary conserved importance of the gene in regulating fatty liver upon nutrition deprivation. Overall, our study identifies a conserved, druggable target to protect the liver from atrophy during starvation.

**One-Sentence Summary:** Cavefish evolved protection from starvation-induced liver damage through reduction of fatty acid uptake regulated by *FATP2*, a mechanism conserved through 400 million years of animal evolution.

## Introduction

Starvation is a severe form of malnutrition that occurs when an individual’s intake of food is inadequate to meet their body’s energy requirements. Prolonged starvation can cause permanent organ damage, stunted growth in children, and eventually death if left untreated (1,2). It is estimated that approximately 10% of the global population suffered from chronic undernourishment in 2021 (3,4), and that approximately 45% of deaths among children under the age of 5 years are linked to undernutrition (5,6). This figure has been steadily increasing in recent years, partly due to factors such as conflicts and climate change (7). Developing interventions aimed at improving starvation resistance is critically needed to fight against nutritional deficiency. Surprisingly, despite the widespread prevalence of starvation, there has been considerably more research focusing on preventing tissue damage resulting from overconsumption than from prolonged hunger.

Notably, the liver’s function and health are compromised during starvation. Almost 90 years ago, the pediatrician Dr. Cicely Williams recognized that chronic severe malnutrition in children leads to fatty liver, and named the condition as ‘kwashiorkor’ (8–11). Subsequently, starvation has been shown to induce accumulation of liver fat (steatosis) in drosophila (12), zebrafish (13,14), and mammals, including mice (15), minks (16), cats (17) & humans (18,19). Notably, cats suffering from sudden anorexia, due to decreased food availability or secondary to a disease, develop feline hepatic lipidosis (FHL), which is the most common hepatobiliary disease in cats. The prognosis from FHL can be positive but only with appropriate and fast nutritional treatment (20) as mortality can be close to 100% without aggressive nutrition therapy (21). Pharmacological interventions to protect the liver during starvation are non-existent.

To uncover potential strategies for protecting the liver, we turned to adaptation strategies observed in nature. Across the animal kingdom, numerous species have developed adaptations to cope with starvation, offering potential perspectives into strategies to combat its detrimental effects (22,23). To gain mechanistic insights, we used a genetically tractable and naturally occurring model of starvation resistance - the *Astyanax mexicanus* model system. Cave populations of this species have adapted to survive in conditions of extreme starvation, while the surface populations of the same species display relatively normal vertebrate physiology. We took advantage of this unique system to study the response of the liver to starvation and identified reduced expression of the fatty acid uptake gene *slc27a2a/fatp2* allowing cavefish to prevent liver damage under starvation. Targeting this pathway in both fish and flies mitigates starvation-induced hepatic steatosis and protects the liver from lipotoxicity in zebrafish. This demonstrates that the identified pathway is evolutionary conserved for over 400 million years, highlighting its potential as a druggable target.

## Results and Discussion

### Higher starvation resistance of cavefish larvae as compared to surface fish larvae

It has been previously shown that cavefish of *Astyanax mexicanus* fare better under starvation compared to their surface cousins when grown under *ad libitum* feeding conditions as they are able to accumulate large fat reserves as adults (24). However, if this ability is independent of feeding and present at earlier developmental stages has not been rigorously addressed. Cavefish, like most fishes, are endowed with nutrition from the mother in the form of yolk. Upon consumption of the yolk, the larvae depend on exogenous nutrients for survival. It is known that cavefish eggs possess approximately 15% larger yolk depots compared to surface fish eggs (25); however, visible yolk regression in cavefish completes by 5.5 days post fertilization (dpf) as compared to 4.5 dpf in surface fish (26), implying that under laboratory conditions *Astyanax mexicanus* larvae enters starvation latest by 6 dpf. To assess if cavefish larvae are more resilient to starvation as compared to surface fish larvae after this time point, we withheld nutrition and tracked daily survival. Surface fish larvae survived robustly until 10 dpf, but afterwards showed a steep reduction in their survival curve (Fig. 1A). In contrast, larvae of two parallelly evolved morphs of cavefish, Pachón and Tinaja, demonstrated more variability in their early survival, in line with previous observations of higher early embryonic mortality in certain laboratory strains; however, once they survived day 6 the cavefish larvae were able to survive up to seven days longer than surface fish without food (Fig. 1A).

**Fig. 1.**
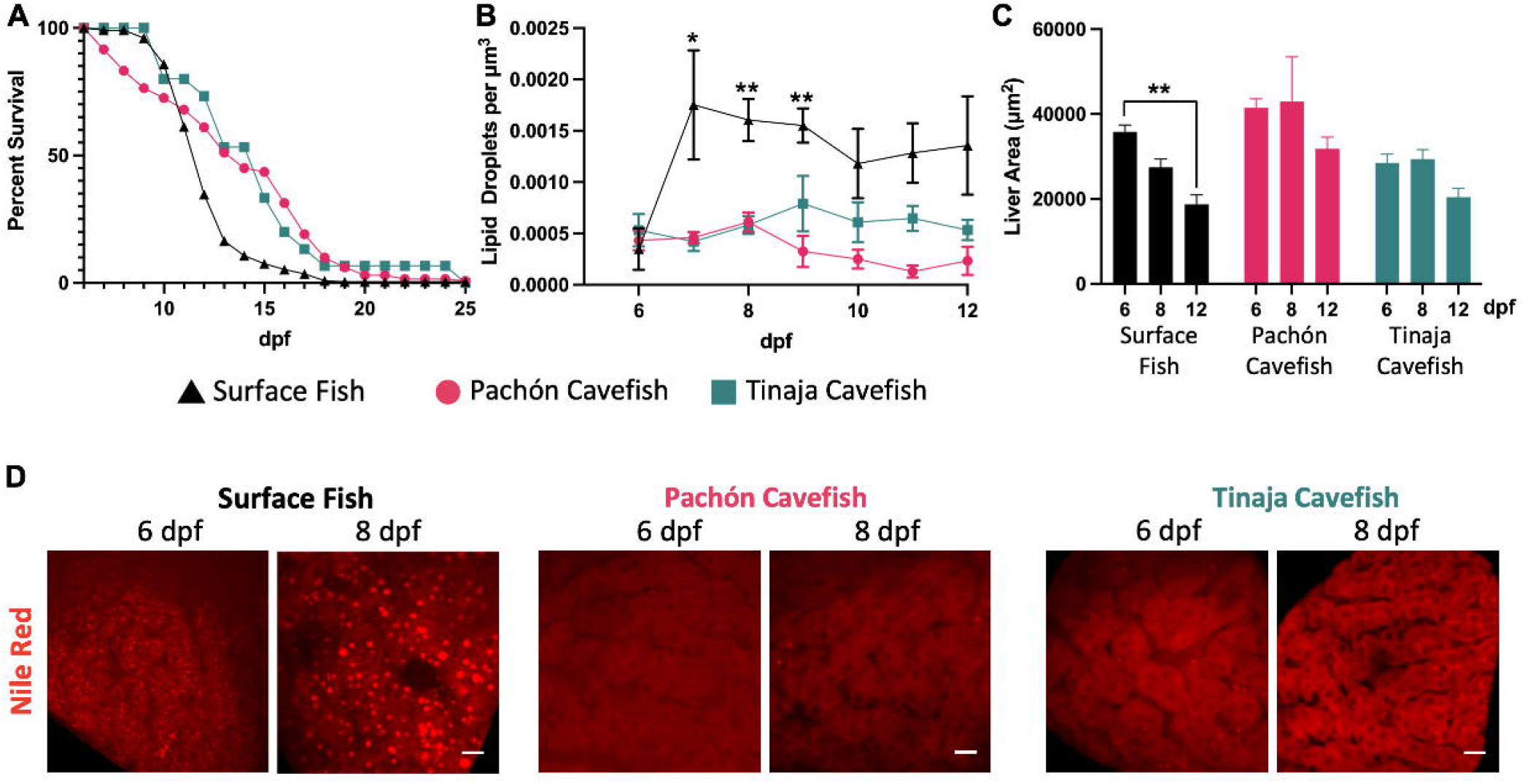
Response of cavefish and surface fish larvae to starvation. **(A)** Survival curve for surface fish, Pachón cavefish and Tinaja cavefish larvae upon starvation are shown by log-rank Kaplan-Meier plots with 95% Confidence Interval. **(B)** Number of hepatic lipid droplets per unit volume from 6 dpf to 12 dpf under fasting. **(C)** Mean ± SEM of liver size in surface fish, Pachón cavefish and Tinaja cavefish at 6 dpf, 8 dpf and 12 dpf without exogenous feeding. ** p-value < 0.01. **(D)** Maximum intensity projections of representative livers from surface fish, Pachón cavefish and Tinaja cavefish stained with Nile Red (red) at 6 dpf and 8 dpf fasting condition. Scale bar = 10 μm

Starvation is known to induce accumulation of liver fat and liver atrophy in vertebrates (15,17). To test if cavefish evolved a different tissue response to starvation, we quantified lipid droplet accumulation in the liver using Nile Red staining. In line with observations from other species, we observed an accumulation of lipid droplets in the liver of surface fish upon starvation, peaking at 7 dpf (Fig. 1B, D). However, strikingly, we did not detect any noticeable increase in lipid droplets in the liver of cavefish during starvation (Fig. 1B, D). Further, we noticed that surface fish display starvation-induced liver atrophy, with the liver size decreasing by 48 % from 6 dpf to 12 dpf (p-value = 0.002) (Fig. 1C). However, the cavefish liver seemed to be more resilient to starvation-induced liver atrophy, with Pachón and Tinaja showing only a more modest reduction of 23 % (p-value = 0.15) and 28 % (p-value = 0.17), respectively, over the six days of starvation. To our knowledge, this is the first report of a model organism that is able to avoid starvation-induced hepatic steatosis.

### Liver atrophy upon accumulation of hepatic lipid droplets

We investigated the relationship between the accumulation of hepatic lipid droplets and liver atrophy upon starvation. It is known from Non-alcoholic Fatty Liver Disease (NAFLD) that liver damage occurs when hepatic steatosis proceeds to steatohepatitis, characterized by the presence of inflammation (27). However, to our knowledge, it is not known if liver damage upon starvation progresses through a similar sequence of events. To investigate this, we turned to the zebrafish as a model system, where we and others have shown robust development of starvation-induced hepatic steatosis (13,14), similar to surface fish. We visualized hepatic lipid droplets in the *Tg(fabp10a:EGFP)* reporter line, which marks hepatocytes with green fluorescence, using Nile Red staining of neutral lipids. We found that at 4 dpf, zebrafish display very little hepatic lipid droplets, while at 6 dpf, without exogenous feeding, zebrafish enter a fasting state with robust hepatic steatosis (Fig. 2A) (13). Following 6 dpf, hepatic lipid droplets decreased (Fig. 2A, C) and liver atrophied (Fig. 2A, G). To quantify macrophage presence as an indicator of inflammation, we took advantage of the double transgenic zebrafish line *Tg(fabp10a:mCherry)*; *Tg(mpeg1*.*1:EGFP)* to visualize hepatocytes and liver macrophages *in vivo*. Notably, due to the pH-stability of mCherry, this combination allowed us to not only visualize macrophages, but also detect the presence of hepatic phagocytic debris in macrophages (Fig. 2B). We found a significant increase in the number of macrophages in the liver at 7 dpf and 8 dpf (Fig. 2B, C), suggesting a transition from steatosis to steatohepatitis.

**Fig. 2.**
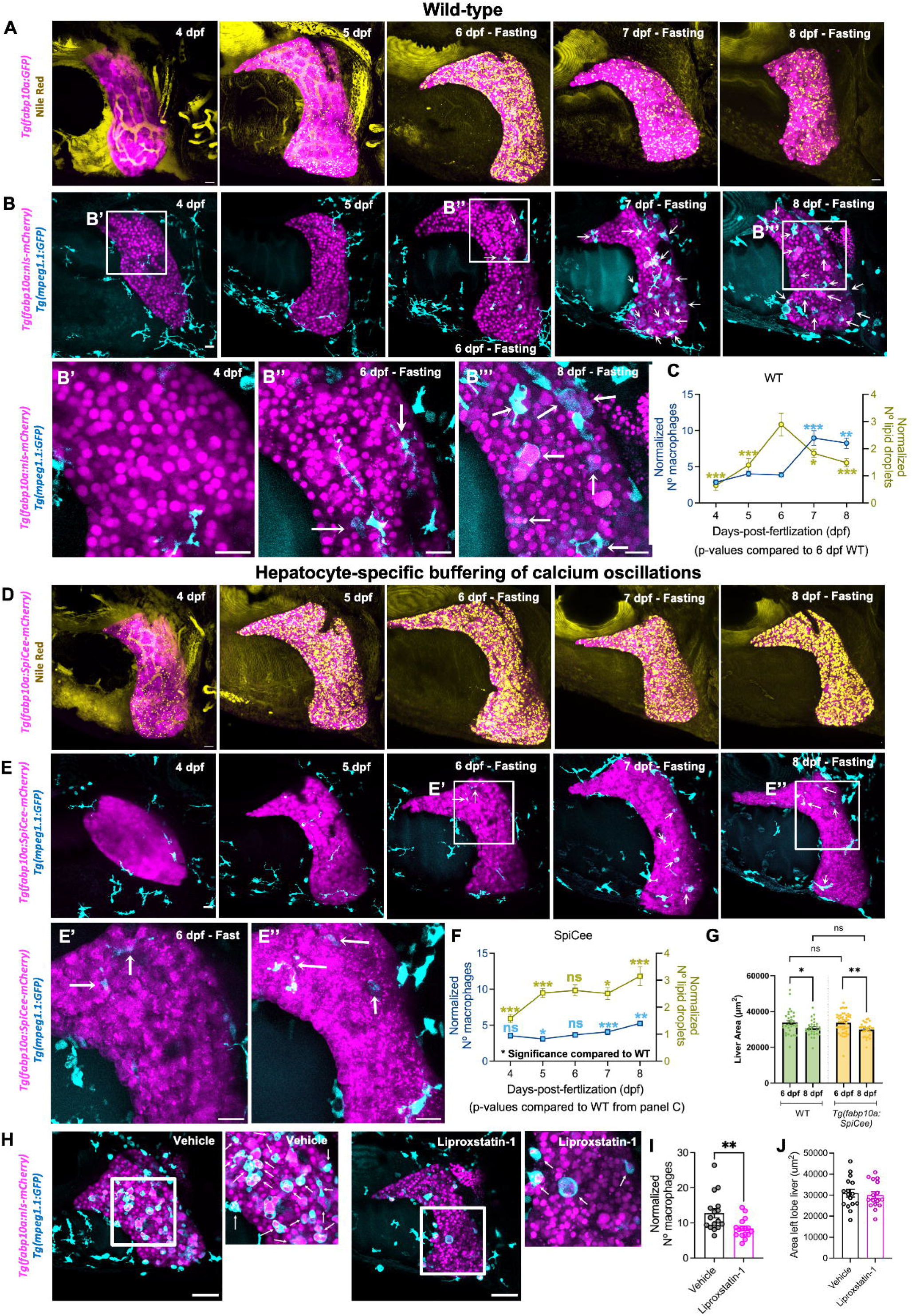
Reduction in the size of liver upon accumulation of lipid droplets during starvation. **(A)** Maximum intensity projections (MIP) of Tg(fabp10a:GFP) (referred as WT) stained with Nile Red. Hepatocytes are false-colored in pink and lipid droplets in yellow. A timeline from 4 dpf to 8 dpf of zebrafish larvae without exogenous food is presented. **(B)** Representative MIP images of timeline from Tg(fabp10a:nls-mCherry); Tg(mpeg1.1:EGFP) zebrafish from 4 dpf to 8 dpf under fasting condition. Arrows label macrophages with hepatocyte debris. A zoom of boxed regions is presented below the images. **(C)** Line trace representing lipid droplets and macrophages normalized to the liver area from 4 dpf to 8 dpf. Comparisons to 6 dpf are represented. p-values: * < 0.05; ** < 0.01, *** < 0.001; ANOVA test followed by post-hoc Tukey test. **(D)** MIP of Nile Red staining of Tg(fabp10a:SpiCee-mCherry), with hepatocytes false-colored in pink and lipid droplets in yellow. Representative images from 4 dpf to 8 dpf fasting are presented. **(E)** Livers from 4 dpf to 8 dpf from fasting Tg(fabp10a:SpiCee-mCherry); Tg(mpeg1.1:EGFP) are presented with MIP. Macrophages with hepatocyte debris are marked with arrows. A zoom of boxed regions is presented below the images. **(F)** Line trace representing lipid droplets and macrophages normalized to the liver area from 4 dpf to 8 dpf in Tg(fabp10a:SpiCee-mCherry) animals. Comparisons to WT are shown. p-values: ns = not significant; * < 0.05; ** < 0.01, *** < 0.001; Student t-test or Mann-Whitney U test depending on normality of the data. **(G)** Mean ± SEM of liver size in wild-type and Tg(fabp10a:SpiCee-mCherry) at 6 dpf and 8 dpf without exogenous feeding. p-values: ns = not significant; * < 0.05; ** < 0.01; Student t-test. **(H)** Snapshots of livers at 8 dpf from Tg(fabp10a:nls-mCherry); Tg(mpeg1.1:GFP) fasting animals treated with 0.17% DMSO (vehicle) or 50 μM liproxstatin-1. Hepatocytes are false-colored in pink and macrophages are false-colored in cyan. White arrows indicate macrophages with hepatocyte phagocytosis. **(I)** Mean ± SEM of the number of macrophages normalized by liver area in vehicle and treated animals. Each point represents a single animal. ** p-value < 0.01, Mann-Whitney U test. **(J)** Mean ± SEM of the liver area in control and liproxstatin-1 treated animals at 8 dpf. Scale bar for all panels = 20 μm.

Next, to investigate the relationship between hepatic lipid utilization and inflammation, we manipulated turnover of lipid droplets in the liver. We had previously shown that buffering of calcium signaling in the hepatocytes, accomplished by hepatocyte-specific expression of a genetically encoded buffer of cytosolic calcium oscillations called *SpiCee* (28), initiates hepatic steatosis at 4 dpf, in the fed state (13). Here, we investigated the hepatic lipid content in the *Tg(fabp10a:SpiCee-mCherry)* animals upon fasting. In 6 dpf animals devoid of exogenous food, hepatocytes from SpiCee-expressing and control animals had similar levels of lipid droplets (compare Fig. 2B, C with Fig. 2D, F). However, while the control animals reduce the droplets, SpiCee-expressing hepatocytes do not show reduction of lipids until 8 dpf, maintaining a steady level of lipid droplets (Fig. 2D, F). Strikingly, livers from SpiCee-expressing animals did not show a significant increase in liver macrophages from 6 – 8 dpf (Fig. 2E, F). This suggests that reducing the turnover of lipid droplets protects the liver from inflammation. However, the liver in SpiCee-expressing animals did undergo atrophy at the same rate as the liver in wild-type animals (Fig. 2G), suggesting that inhibiting lipid utilization cannot protect the liver from atrophy.

In accordance with the model where the lipid droplet turnover correlates with inflammation, several studies have suggested the role of lipid droplets as a store of potential pro-inflammatory molecules (29–31). Degradation of lipid droplets releases polyunsaturated fatty acids (PUFA) that can react with reactive oxygen species (ROS) to generate cytotoxic lipid peroxidation molecules (32). To evaluate whether lipid peroxides generated by lipid droplet degradation modulates the inflammatory response in the zebrafish liver, we treated *Tg(fabp10a:nls-mCherry);Tg(mpeg1*.*1:GFP)* animals with liproxstatin-1 (lipid peroxidation inhibitor) from 6 dpf to 8 dpf under starvation. Reduction of lipid peroxidation decreased macrophages in the liver (Fig. 2H, I), suggesting a role of lipids in steatohepatitis. However, liproxstatin-1 treatment could not protect the liver from atrophy, as the size of the liver between control and liproxstatin-1 treated animals was the same at 8 dpf (Fig. 2J).

Overall, our data provides evidence that reduction of lipid flux or inflammation does not protect the liver from atrophy during starvation in zebrafish larval stage.

### Inhibition of hepatic lipid accumulation protects the liver from atrophy during starvation

As the liver size reduces after accumulation of lipid droplets, irrespective of lipid flux or inflammatory status, the best strategy to protect the liver is potentially to avoid lipid droplet formation during starvation, as done by cavefish. With this, potentially cytotoxic lipid products are not accumulated in the hepatocytes. To understand how cavefish prevent the formation of lipid droplets upon starvation, we compared transcriptomes of livers from fed and fasting state. We profiled the liver from 4 dpf (fed) and 6 dpf (fasting) zebrafish larvae, and 5 dpf (fed) and 7 dpf (fasting) surface fish and Tinaja cavefish larvae. Upon performing 2-D dimensionality reduction of the transcriptomes using multidimensional scaling (MDS), we observed that the species was separated on the X-axis, suggesting species-specific responses to starvation (Fig. 3A). However, the Y-axis represented the fed-to-fast transition for the two species, suggesting a shared response to nutrition deprivation (Fig. 3A). Notably, this underscores the fact the cavefish larvae also undergo starvation after 6 dpf.

**Fig. 3.**
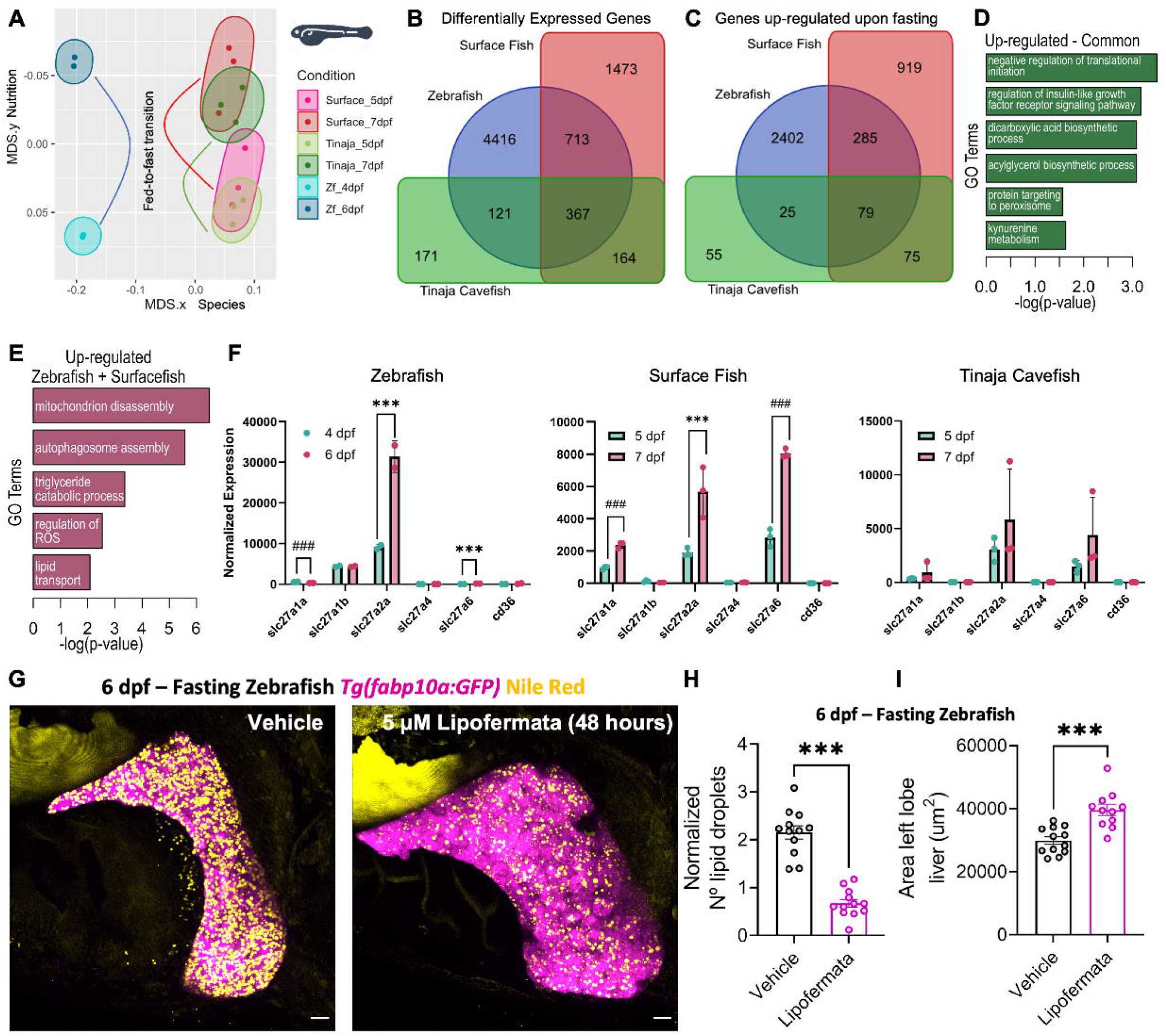
*Slc27a2a* is responsible for starvation-induced lipid accumulation. **(A)** Multidimensional Scaling (MDS) plot of gene expression changes in the liver upon fasting in zebrafish, surface fish and Tinaja cavefish. For zebrafish, livers from 4 dpf were compared to 6 dpf, while for surface and cavefish the comparison was made between 5 dpf and 7 dpf. **(B, C)** Venn-diagram of differentially expressed genes (DGE) (B) and genes up-regulated by fasting (C) for the three animals. **(D, E)** Gene ontology (GO) analysis for genes up-regulated by fasting in all the three animals (D) and for zebrafish and surface fish only (E). **(F)** Barplot displaying the changes in lipid transporters upon fasting. *** False-discovery rate (FDR) < 0.05 and log2(fold-change) > 1.5. ### (FDR) < 0.05, but log2(fold-change) < 1.5. **(G)** Maximum intensity projections of 6 dpf fasting *Tg(fabp10a:GFP)* (pink) with Nile Red staining (yellow) treated with 5 μM of lipofermata or 0.01% of DMSO (vehicle) from 4 dpf to 6 dpf fast. Scale bar = 20 μm. **(H, I)** Bar plot with mean ± SEM of the number of lipid droplets per liver (H) and liver size (I) in vehicle and lipofermata treated animals. Each point represents a single animal. ns = not significant; *** p-value < 0.001, student t-test.

Differential gene expression (DGE) analysis identified 367 genes to be modulated in all three groups (Fig. 3B). Amongst these, 79 genes were up-regulated in the starvation state (Fig. 3C). Gene ontology (GO) analysis revealed that these 79 genes were enriched in “regulation of translational initiation”, “insulin-like growth factor signaling”, “acylglyceroal biosynthetic pathway” and “protein targeting to peroxisomes” (Fig. 3D). We next focused on genes up-regulated upon starvation in zebrafish and surface fish, but not in cavefish. These 285 genes were enriched in “mitochondrion disassembly”, “autophagosome assembly”, “triglyceride catabolic process”, “regulation of ROS” and “lipid transport” (Fig. 3E). Among the lipid transporters, we observed that the long-chain fatty acid transporter, *slc27a2a*, was differentially regulated in zebrafish and surface fish, but not in cavefish (Fig. 3F).

The lipid transporter *slc27a2a* is the fish homologue of SLC27A2, also known as FATP2 (fatty acid transport protein 2), and belongs to the SLC27 family of lipid transporters (33).

To evaluate the role of *slc27a2a* during starvation, we again turned to the zebrafish model and treated zebrafish larvae with lipofermata, a specific inhibitor for SLC27A2 (34), from 4 dpf to 6 dpf, and checked the impact of the drug at 6 dpf, and for two days post-treatment (dpt). At 6 dpf (0 dpt), livers from lipofermata-treated starved animals showed a dramatic reduction of the number of lipid droplets accumulated within the hepatocytes compared to vehicle-treated animals (Fig. 3G, H). At 8 dpf (2 dpt), we observed an accumulation of lipid droplets in the liver, suggesting the inhibition is reversible (Fig. S1).

Next, we quantified liver size under the same treatment regimen and observed a striking increase in the size of the liver (Fig. 3I). Notably, the increase in liver size was present at 6 dpf (0 dpt) and persisted at-least until 8 dpf (2 dpt) (Fig. S1).

Our experiments suggest that inhibition of fatty acid uptake can reduce accumulation of lipid droplets in the liver, which in turn can reduce liver atrophy upon starvation.

### SLC27A2 / FATP2 is an evolutionary conserved regulator of starvation-induced hepatic steatosis

To test the evolutionary conservation of SLC27A2 / FATP2 in starvation-induced steatosis, we turned to drosophila, which shows robust accumulation of lipid droplets in oenocytes, the evolutionary homologues of hepatocytes in the fly (12). Using tissue-specific RNA interference (RNAi), we knocked down the drosophila homologue of FATP2. Two independent siRNA lines were used. Upon starvation, the oenocytes from control animals displayed an accumulation of lipid droplets (Fig. 4A, B). However, RNAi-mediated knock-down of FATP2 significantly reduced the number of lipid droplets (Fig. 4A, B).

**Fig. 4.**
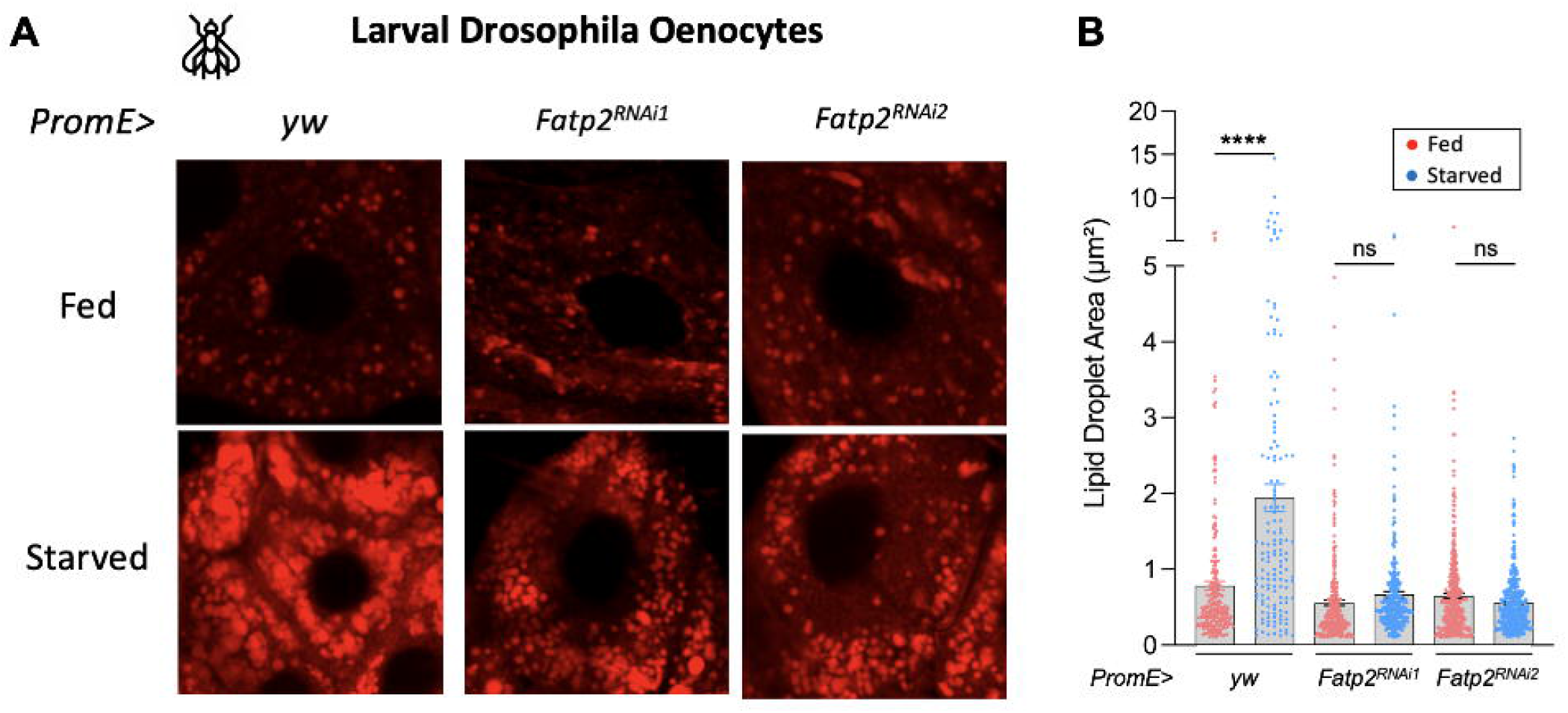
FATP2 is an evolutionarily conserved regulator of starvation-induced lipidosis. **(A)** Confocal images of drosophila oenocytes from fed and starved larvae. Control or oenocyte-specific FATP2 RNAi were evaluated. **(B)** Barplot comparing lipid droplet area in fed or fasting condition in control and FATP2 RNAi animals. ns > 0.05, **** p-value < 0.0001, ANOVA test followed by post-hoc Tukey test.

This suggests that the importance of SLC27A2 / FATP2 in regulating starvation-induced hepatic steatosis is conserved across approximately 400-million years of evolution (35).

However, the accumulation of lipid droplets can severely compromise the health of the liver. Our study shows that the optimal solution to starvation-induced liver injury, identified by naturally starvation-resistant cavefish, is to prevent the accumulation of hepatic lipid droplets. In future, it would be of interest to understand the upstream regulators of slc27a2a gene expression to uncover the molecular link between lack of nutrients and lipid uptake in hepatocytes. This could provide additional targets to protect the liver from atrophy during starvation.

## Material and Methods

### *Astyanax mexicanus* husbandry

Surface, Tinaja and Pachón morphs of *Astyanax mexicanus* were raised in polycarbonate recirculating aquaculture racks (Pentair Aquatic Eco-Systems, Apopka, FL), with a 14:10 h light:dark photoperiod. Each rack system is equipped with mechanical, chemical, and biological filtration and UV disinfection. Water quality parameters are monitored daily as described in previous studies (36). Fish were housed at a density of ∼2 fish per liter. Fish embryos spawned at the same time from different parent tanks were mixed in order to reduce effects from specific backgrounds. Only embryos screened as healthy were raised and randomly selected for experiment. Embryos up to 5□days after fertilization were maintained at 24.5□°C in E2 embryo media. Animal husbandry followed protocol 2021-122 approved by the Institutional Animal Care and Use Committee (IACUC) of the Stowers Institute for Medical Research. Housing conditions meet federal regulations and are accredited by AAALAC International.

### Survival curve for *Astyanax mexicanus*

At 5 dpf, larvae were randomly divided into two groups. Each group included ten biological replicates, and 20 embryos as one biological sample. One group, as the controls, were fed with *Artemia nauplii* (Brine Shrimp Direct, Ogden, Utah) once per day. The other, as the starvation group, were not fed. The fish were identically treated in all other aspects, and the number of living fish was recorded daily. Dead fish were removed each day. The experiment was repeated 3 times independently.

### Nile Red staining, confocal imaging, and analysis for *Astyanax mexicanus*

Nile Red staining was performed as previously described in (36) with some modifications. Briefly, stock solution of Nile Red (Invitrogen; # N-1142) was made by dissolving in acetone to a concentration of 500ug/mL and stored in the dark at 4°C. A final working solution, diluted to 1/300 in embryonic E2 media, was prepared. For the staining procedure, 10 larvae per well were incubated in 12-well cell culture plates (Corning; # CLS3513) containing 2□mL in the working solution for 1h at 25 °C and protected from light. Following incubation, stained larvae were euthanized with 500□mg/L MS-222, washed with PBS, and then fixed in 4% paraformaldehyde for 20 minutes at room temperature prior to imaging. Samples were placed on a glass-bottom FluoroDish, and images captured using a Nikon Eclipse Ti2 inverted microscope equipped with a Yokagawa Spinning Disk Confocal with a 40x/1.15 NA objective. Nile Red was excited with a 561nm laser and emission was collected between 579-631 nm. Z-stack images were acquired with 2048x2044 pixel size and step size of 1 μm.

Image quantification was performed in Imaris (Oxford Instruments). We first manually segmented the liver to find the volume and mask the lipid fluorescence channel. Lipid droplets were created using the spots function set to 0.8μm seed size. Liver area was measured in ImageJ (37) by outlining the whole liver in a maximum projected image with the polygon selection tool.

### RNA-Sequencing of *Astyanax mexicanus* livers

For RNA-Seq, RNA was isolated from livers at the corresponding stages. For this, livers from 40 larvae / replicate were dissected and collected in the lysis buffer from ReliaPrep™ RNA Miniprep Systems (Promega, Z6110). RNA was isolated from the lysed tissue by following the manufacture’s instruction. cDNA synthesis, library preparation and Illumina sequencing was performed according to the manufacturer’s instructions using the TruSeq Stranded mRNA Preparation Kit for Illumina (Ilumina #20020594). Purified libraries were quantified using a Quibit fluorometer (Life Technologies) and the quality assessed on the Agilent Bioanalyzer (Agilent Technologies). Sequencing was performed on the Illumina NextSeq 500 instrument as High Output with single-end 75bp reads. Following sequencing, raw reads were demultiplexed into Fastq format using Ilumina bclfastq2 v2.18.

### Mapping of transcriptome from *Astyanax mexicanus* livers and comparison to zebrafish transcriptome

Raw reads were mapped using STAR aligner v2.7.3a (38) against the UCSC genome astMex_2.0 genome, using Ensembl 102 gene models for annotations. Transcript abundance was quantified using RSEM v1.3.0. For normalization, differential gene expression analysis, and gene ontology (GO analysis), iDEP version 0.951 (39) was utilized with default parameters. For differential gene expression, a fold-change of 2 and false-discovery rate of 0.1 was used for cut-off. Data for zebrafish liver transcriptome from fed and fasting condition was taken from our previous publication (13). Common and uniquely differentially expressed genes were compared using an online Venn Diagram tool (40).

### Zebrafish strains and husbandry

Wild-type or transgenic zebrafish of the outbred AB strain were used in all experiments. Zebrafish were raised under standard conditions at 28 °C. Animals were chosen at random for all experiments. Zebrafish husbandry and experiments with all transgenic lines will be performed under standard conditions in accordance with institutional (Université Libre de Bruxelles (ULB)) and national ethical and animal welfare guidelines and regulation, which were approved by the ethical committee for animal welfare (CEBEA) from the Université Libre de Bruxelles (protocols 854N-865N). The following transgenic zebrafish strains were used in this study: *Tg(fabp10a:SpiCee-mCherry)*^*ulb16*^ (13), *Tg(mpeg1*.*1:EGFP)*^*gl22*^ (41), *Tg(fabp10a:nls-mCherry)*^*mss4*^ (42), *Tg(fabp10a:EGFP)*^*as3*^ (43). The developmental stages of zebrafish used in experiments are prior to sex specification. All zebrafish were healthy and not involved in previous procedures.

### Pharmacological treatments

Stocks for Lipofermata (MedChem, HY-116788) and Liproxstatin-1 (Sigma, SML1414) were prepared in DMSO and stored at -80ºC. Working concentration for Lipofermata was 5 μM and for Liproxstatin-1 was 50 μM. Incubation was performed in a petri dish sealed with parafilm to avoid evaporation. The petri dish was kept in dark during the treatment. DMSO was used as a control.

### Imaging of the zebrafish liver macrophages

Animals were anesthetized in 0.02% tricaine methanesulfonate (MS-222; E10521; Sigma-Aldrich, Darmstadt, Germany) and mounted in 1% Low-Melt Agarose containing 0.02% Tricaine MS-222 (50080; Lonza, Basel, Switzerland) and imaged on a glass-bottomed FluoroDish (FD3510-100; World Precision Instruments (WPI), Sarasota, Florida) using a Zeiss LSM 780 confocal microscope. Livers were imaged using a 40x/1.1N.A. water correction lens. Samples were excited with 488 nm for *Tg(mpeg1*.*1:EGFP)* and 543 nm for *Tg(fabp10a:nls-mCherry)* or *Tg(fabp10a:SpiCee-mCherry)*. The imaging frame was set at 1024x1024 pixels, and the distance between confocal planes was set at 3 μm for Z-stack to cover, on average, a thickness of 100 μm. Analysis was performed in Fiji (37). Macrophages in contact with hepatocytes were counted manually. Liver area was calculated by outlining the region using the magic wand area selection tool of a Z-Stack projection, followed by the “Analyze” – “Measure” (Area) command. Normalized number of macrophages was calculated using the next formula:

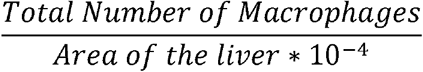

### Nile Red staining, confocal imaging, and analysis for zebrafish liver

Imaging of the lipid droplets in the zebrafish liver was performed exactly as mentioned in our previous publication (13), which includes the time-line of hepatic lipid droplets in wild-type condition from 4 – 6 dpf. In brief, Nile Red stock was prepared by dissolving 500 μg/ml of Nile Red (Art. No. 7726.2; Carl Roth, Germany) in acetone and filtering the solution with a 0.2-μm filter. Stock solution was stored at 4ºC in the dark. On the day of imaging, a fresh working solution of 1/300 in Embryonic medium E2 was prepared. The larvae were incubated in the working solution for 1 h at 28ºC, protected from light. After incubation, larvae were washed once with E2 medium. Animals were anesthetized with 0.02% tricaine MS-222, mounted in 1% Low-Melt Agarose containing 0.02% of tricaine MS-222, and imaged on a glass-bottomed FluoroDish using a Zeiss LSM 7890 confocal microscope. Livers were imaged using a 40x/1.1N.A. water correction lens. The imaging frame was set at 1024 x 1024 pixels, and the distance between confocal planes was set at 2 μm for Z-stack to cover, on average, a thickness of ∼100 μm. Samples were excited with 488 nm, and a spectral detector was used to collect fluorescence corresponding to EGFP (500 – 508 nm) and neutral lipids labeled by Nile Red (571 – 579 nm).

Analysis was performed in Fiji (37). Lipid droplets were outlined manually with a circular selection tool. Only lipid droplets (marked with red fluorescence) present within hepatocytes (marked with green fluorescence) were counted. Area of the hepatocyte region was calculated by outlining the region using the magic wand area selection tool, followed by the “Analyze” – “Measure” (Area) command.

### Drosophila stocks and husbandry

Flies were maintained at 25°C, 60% relative humidity, and 12 h light/dark cycles. Larvae were reared on the standard cornmeal and yeast-based diet (0.8% cornmeal, 10% sugar, and 2.5% yeast. For starvation treatment, early L3 larvae (∼72 hours after egg laying) were kept with Kimwipe soaked with 1x Phosphate Buffered Saline for 16∼18 hours. Fly stocks used in this study were as follows: PromE-Gal4 (BDSC #65405), Fatp2 RNAi (#1-BDSC #55271, #2-BDSC #44658). ywR flies (a gift from Eric Rulifson) were crossed to PromE-gal4 as control.

### Nile Red staining, confocal imaging, and analysis for Drosophila oenocytes

Larva was dissected in Schneider’s Drosophila medium (ThermoFisher #217-20024) (normal fed larvae), or in 1x Phosphate Buffered Saline (1x PBS) (Starved larvae). The dissected larvae were fixed with 4% paraformaldehyde (Electron Microscopy Sciences #15710) for 20 min at room temperature. The stock Nile Red was prepared by dissolving 1mg of Nile Red (Invitrogen™ #N1142) In 1ml of dimethyl sulfoxide (DMSO). The larval tissues were incubated in 2ug/ml Nile Red working solution for 1 hour at room temperature, protected from light. The samples were then washed and mounted on the slides using ProLong Diamond Antifade Mountant (Thermo Fisher #P36961). The lipid droplets were imaged with an FV3000 Confocal Laser Scanning Microscope (Olympus). The imaging frame was set at 1024 x 1024 pixels. Samples were excited with 555nm, the range of 580nm-610nm was used to detect the Nile Red signal.

The images were analyzed by Olympus cellSens Dimensions software. The lipid droplets were identified by using the region of interest (ROI) and adjusting the fluorescent threshold to include the lipid droplets inside the oenocytes. The area is calculated automatically by Olympus cellSens Dimensions software after identifying the lipid droplets.

### Statistical analysis

Statistical analysis was performed using GraphPad Prism software (version 9.3.1; GraphPad Software, San Diego, CA). The test used for comparison is mentioned in the figure legends of respective graphs. For all analysis, data was tested for outliers and normal distribution. For non-normal distributed data (or if the test for normal distribution could not be performed because of small sample size), a nonparametric test was used. No data were excluded from analysis. Blinding was not performed during analysis.

## Acknowledgments

We thank the members of IRIBHM fish facility and M. Martens and J.-M. Vanderwinden from the Light Microscopy Facility for technical assistance at ULB. We are grateful to the cavefish facility at the Stowers Institute for Medical Research cavefish husbandry support cavefish. Authors would also like to thank Amanda Lawlor from Stowers Sequencing and Discovery Genomics team for help regarding Library Preparation and sequencing of cavefish samples. We thank Bloomington Drosophila Stock Center (supported by NIH P40OD018537) for fly stocks. We thank Eric Rulifson for sharing the yw fly stock.

## Funding

Fonds de la Recherche Scientifique (FNRS) grant 40005588

(SPS) Fonds de la Recherche Scientifique (FNRS) grant 40013427 (SPS)

Stowers Institute grants (NR)

National Institute of Health (NIH) 1DP2AG071466-01 (NR)

National Institute of Health (NIH) R24OD030214 (NR)

National Institute of Health (NIH) R01AG058741 (HB)

National Institute of Health (NIH) R01AG075156 (HB)

National Science Foundation (NSF) CAREER 2046984 (HB)

## Author contributions

Conceptualization: NR, SPS Methodology: MPM, AEC, CC, PL

Investigation: MPM, AEC, CC, PL, CP, AL, FL

Visualization: MPM, AEC, CC, MCM, PL

Funding acquisition: NR, SPS

Project administration: SPS

Supervision: HB, NR, SPS

Writing – original draft: NR, SPS

Writing – review & editing: AEC, HB, NR, SPS

## Data and materials availability

All data needed to evaluate the conclusions in the paper are present in the paper and/or the Supplementary Materials. Additional data related to this paper may be requested from the authors. The sequencing data are available at NCBI Gene Expression Omnibus (GEO) under GSE244648 with the review token mvkjqsyqbzknzmf.

