## Supplementary Figure 1 for "Starvation resistant cavefish reveal conserved mechanisms of starvation-induced hepatic lipotoxicity"

† These authors contributed equally.

**Table of Content**

Figure S1: Enhancement of liver size for two days after removal of lipofermata treatment. 3

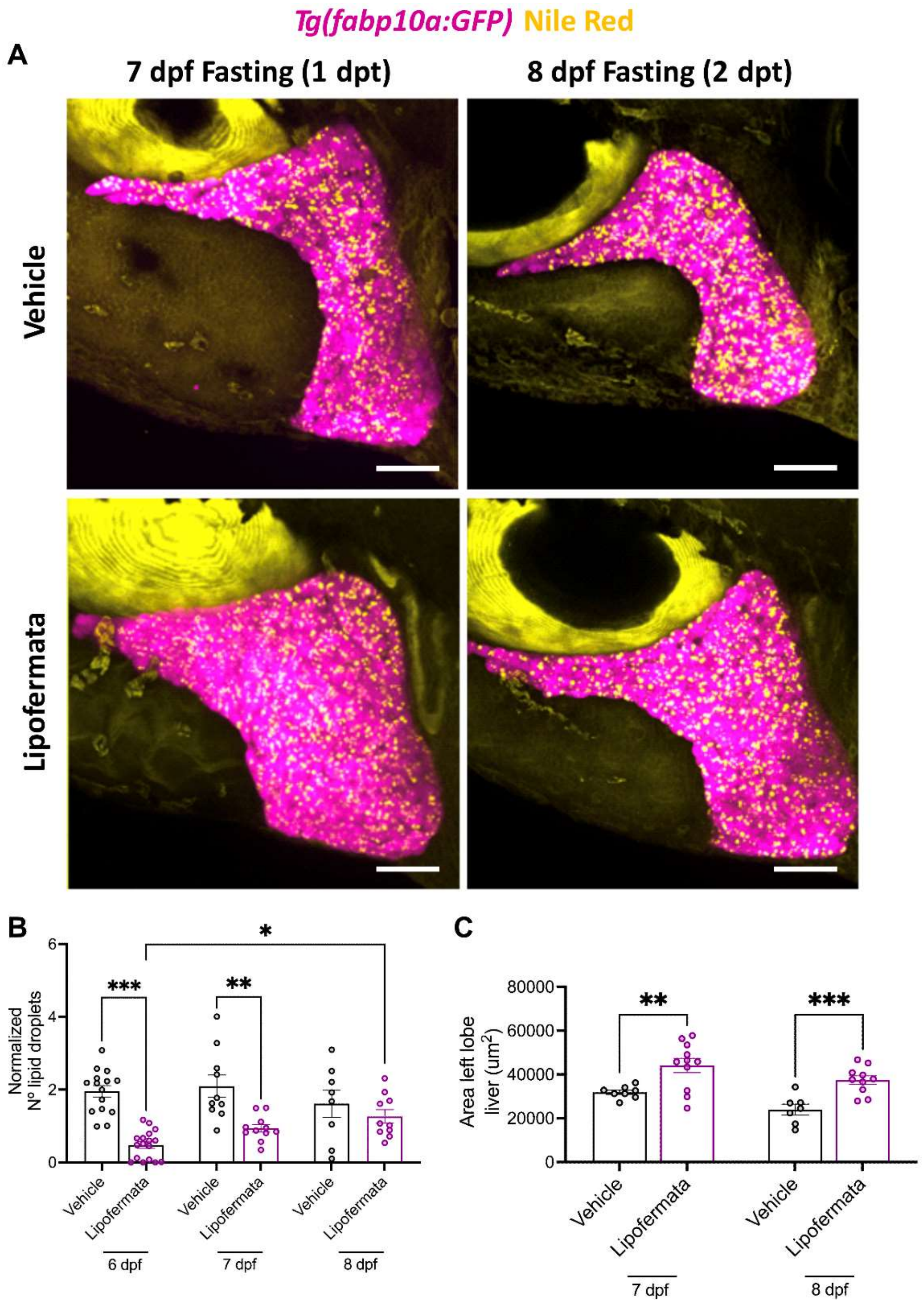


**Figure S1: Enhancement of liver size for two days after removal of lipofermata treatment.** *Tg(fabp10a:GFP)* animals were treated with 5 µM of lipofermata or 0.01% of DMSO (vehicle) from 4 dpf to 6 dpf fast. The drugs were washed and the animals were allowed to recover. **(A)** Maximum intensity projections of 7 dpf and 8 dpf fasting *Tg(fabp10a:GFP)* (pink) with Nile Red staining (yellow) treated with 5 µM of lipofermata or 0.01% of DMSO (vehicle) from 4 dpf to 6 dpf fast. Scale bar = 20 µm. **(B, C)** Bar plot with mean ± SEM of the number of lipid droplets per liver (B and liver size (C) in vehicle and lipofermata treated animals. Each point represents a single animal. * p-value < 0.05, ** p-value < 0.01, *** p-value < 0.001, student t-test.
